# Endotoxin, unidentified material, and low amounts of defined toxicants are confounding factors when using the air pollution Standard Reference Material (SRM) 2786 in biological experiments

**DOI:** 10.1101/2023.08.18.553903

**Authors:** Charles Miller

**Affiliations:** Department of Environmental Health Sciences and the Tulane Cancer Center Tulane University School of Public Health and Tropical Medicine, New Orleans, LA 70112, USA

## Abstract

The National Institute of Standards and Technology (NIST) provides the reference sample SRM 2786 for use in analytical studies of particulate air pollution. The sample originated from an atypical source and possibly contains materials that are unrelated to air pollution. Investigators conducting studies with SRM 2786 should note the relatively low amounts of toxic components listed in the certificate of analysis as well as a large amount of uncharacterized material. Additionally, there is no unsized reference material that can be used as a control for the particulate nature of SRM 2786. The presence of a low amount of endotoxin in SRM 2786 is reported here. Endotoxin and other uncharacterized material in SRM 2786 may influence results and conclusions for biological studies using this reference material.

## Introduction

Airborne particulate matter (PM) is complex and variable in its composition and has been linked to adverse health effects (1). Consequently, particulate matter of 10 microns (PM10) and 2.5 microns (PM 2.5) in the air are monitored to protect health. The US EPA and the World Health Organization have established standards of 12 ug/m^3^ and 5 ug/m^3^, respectively, as acceptably safe levels of PM2.5 in air (2, 3). Airborne PM is regulated by its size rather than its composition and toxicological properties. For example, PM2.5 in the air might come from a source such as engine exhaust, or natural sources such as wind-blown soils. Biological agents such as bacteria and viruses may be present in particulate matter. Particulates from these different sources produce different effects when inhaled, but they all are considered PM2.5 for the purpose of regulation.

A particulate matter sample, SRM 2786, was prepared and partially characterized by the National Institute of Standards and Technology (NIST) of the USA (4). This material was intended to represent urban PM2.5 air pollution. But rather than being directly isolated on standard air sampling media, this material was obtained from “an air intake filtration system of a major exhibition center in Prague, Czech Republic.” This source of material is problematic because the air filter type, usage time, and amount of air filtered are unspecified. Commercial air filters that are in place for extended periods of time can be contaminated with non-airborne material and often provide a source for microbial growth. Thus, SRM 2786 may not accurately represent urban PM2.5 air pollution. While it may serve as an appropriate source for analytical studies, SRM 2786 was not specifically prepared or characterized for use in biological or toxicological experiments.

The material from the commercial air filter was fractionated by size to < 4 um and then partially characterized for numerous elements and organic molecules as described in its certificate of analysis (4). A summary of these analytical categories and their amounts are shown in Table 1 below. Low amounts of potentially toxic metals and organics were measured in this sample. Seventy-eight percent of the sample (by weight) is unknown, and about 20% consists of elements (mostly sodium, magnesium, calcium, iron, aluminum, and chlorine). Some of the metal components of SRM 2786 will not remain PM2.5 when placed in aqueous or biological environments since they likely exist as soluble salts. Additionally, SRM 2786 was characterized for 51 specific polycyclic aromatic hydrocarbons (PAHs), 7 nitro-PAHs (N-PAHs), 6 polybrominated diphenyl ethers (PBDEs), 17 chlorinated dioxins and dibenzofurans (Dioxins), 3 hexabromocyclododecane isomers (HBCDs), 3 anhydrosugars (levoglucosan, mannosan, glactosan) and moisture content. The polycyclic aromatics and halogenated organic compounds collectively constitute <0.1% of SRM 2786.

**Table 1.**
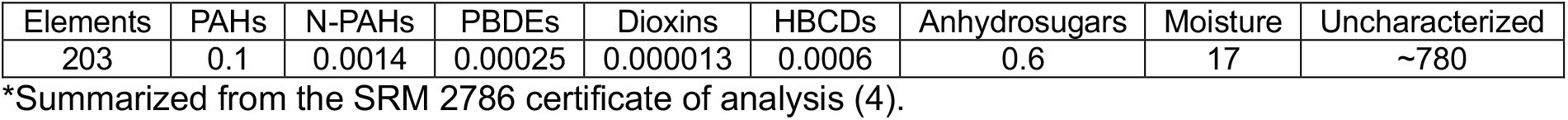
Summary of the analyte groups in SRM 2786 (g/kg)*.

| Elements | PAHs | N-PAHs | PBDEs | Dioxins | HBCDs | Anhydrosugars | Moisture | Uncharacterized |
| --- | --- | --- | --- | --- | --- | --- | --- | --- |
| 203 | 0.1 | 0.0014 | 0.00025 | 0.000013 | 0.0006 | 0.6 | 17 | ~780 |

\*Summarized from the SRM 2786 certificate of analysis (4).

Hill, et al used SRM 2786 as a representative PM2.5 urban air pollution source to induce inflammation and tumor promotion in a murine model of lung cancer (5). However, the uncharacterized material and other variables mentioned above leave questions regarding the bioactive components of SRM 2786. One possibility is that the bioactive component of SRM 2786 is biological rather than anthropogenic in nature.

### Research methods and results

Endotoxin is a potent, natural, and well-known initiator of inflammation (6). Due to the atypical origin of SRM 2786 described above, endotoxin is a candidate for its inflammatory effects. To test for the presence of endotoxin, one ml of sterile water was added to approximately 100 mg SRM 2786 to produce a final concentration of 100 mg/ml. The sample was vortexed extensively to dissolve as much of the material as possible. A Pierce chromogenic endotoxin test kit (ThermoFisher A39552) was used as directed to detect the presence of endotoxin. After the reaction was stopped, the assay plate was allowed to stand for 1 hour and then 150 ul of the assay supernatant was pipetted into a new 96-well plate. This transfer eliminated the interfering sedimented material at the bottom of the original assay plate. The SRM 2786 sample was incubated with the chromogenic substrate alone (no *Limulus* enzyme) to verify that there was no spontaneous activity. Spectrophotometric absorbance readings were taken on the wells of the new plate at 405 nm to detect nitroanilide, the indirect product of the bioassay that indicated the presence of endotoxin. Known amounts of endotoxin were compared to different concentrations of SRM 2786 to determine the amount of endotoxin present (Table 2, Figure 1).

**Table 2.** Endotoxin assay and levels in SRM 2786.

| Endotoxin units,<br>standard curve | Abs. 405 nm (+/- S.D.) | SRM 2786<br>(ug) | Abs. 405 nm (+/- S.D.) | Endotoxin<br>(units/ug SRM 2786) |
| --- | --- | --- | --- | --- |
| 1 | 0.95 (0.05) | 50 | Out of range | - |
| 0.33 | 0.78 (0.11) | 16.7 | 0.94 (0.04) | ~ 0.06 |
| 0.1 | 0.54 (0.05) | 5.5 | 0.74 (0.03) | ~ 0.06 |
| 0 | 0<br>(background<br>subtracted) | - | - | - |

**Figure 1.**
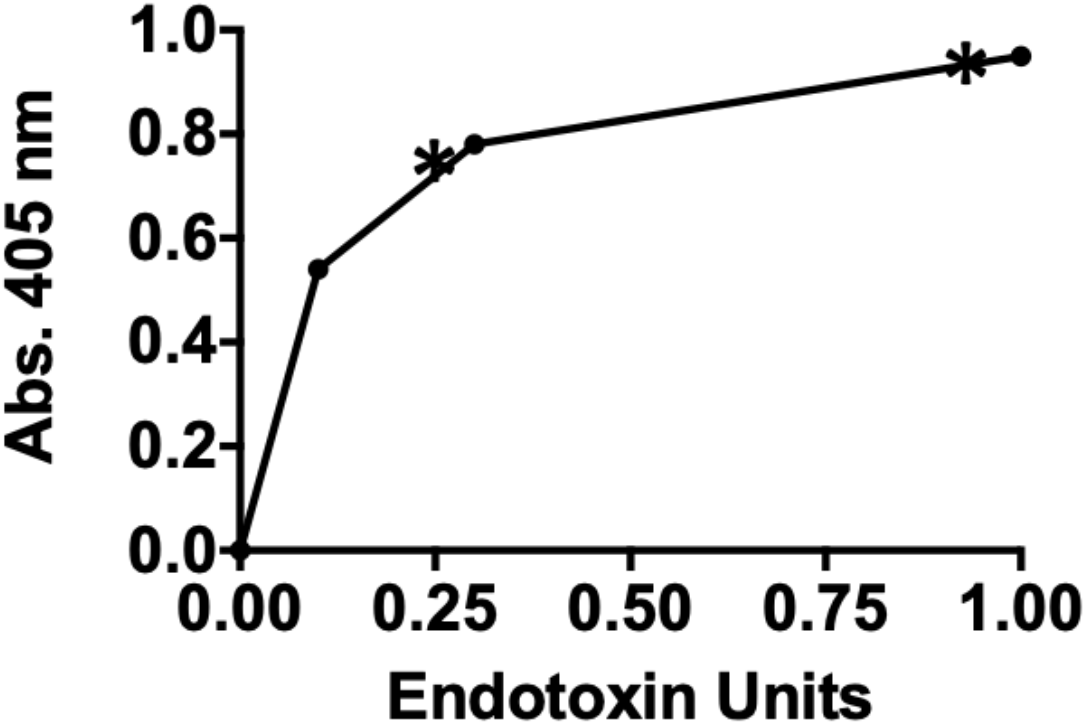
Data plotted from the endotoxin assay. The standard curve is reflected by the connected circular points. Two dilutions of SRM 2786 (16.7 and 5 ug) are indicated by asterisk (*) symbols. Interpolation from the intercepts on the X axis indicate the amount of endotoxin in the samples. The absorbance at 405 nm reflects the nitroanilide product from the indirect assay for endotoxin.

Extrapolations from the endotoxin standard curve indicated SRM 2786 sample contains approximately 0.06 endotoxin units per ug of material. Based on previous studies, it is unknown if this small amount of endotoxin is sufficient to cause inflammatory responses and cytokine expression in experimental systems (7-8). It is important to note that endotoxin can behave synergistically with other airborne pollutants (9). The SRM2786 sample was also tested for viable microorganisms. Five mg SRM 2786 in water was spread onto tryptic soy agar plates and compared with a vehicle control (water alone). Plates were incubated at 22 and 37 °C for 5 days. No microbial growth was detected from the SRM 2786 material. Thus, SRM 2786 may lack viable microbes, but it does contain endotoxin. Endotoxin can be present in the air as a natural microbial product or may have come from microbial growth on the commercial air filter that provided the starting material for SRM 2786. The latter possibility becomes probable if the commercial air filter source was used for an extended period with sufficient humidity and other factors that favored microbial growth.

## Summary

SRM 2786 is a complex reference material that was partially characterized for selected chemical components and size. It is suitable as a reference standard for analytical studies of PM2.5 but should be used with caution in biological systems since it contains low amounts of endotoxin and considerable amounts of uncharacterized material. No experiments have been performed to discern whether the particulate nature of SRM 2786 is required for its biological effects and there is no unsized complementary starting material for comparison. Other common components of PM 2.5 such as black carbon, sulfate, nitrate content, etc., as well as the mutagenicity of this reference sample, are unknown. Environmental studies of PM2.5 tend to focus on anthropogenic air pollution sources which are regulatable. As levels of anthropogenic PM2.5 decline due to the reduced use of fossil fuels, natural components will represent a greater proportion of PM2.5 in the air. Endotoxin, microbes, and other natural materials can be abundant bioactive components of PM2.5 and can produce strong inflammatory responses (6-10). Natural as well as anthropogenic particulate matter should be considered when conducting studies on the health effects of air pollution.

